# Interneuron origins in the embryonic porcine medial ganglionic eminence

**DOI:** 10.1101/2020.12.18.423526

**Authors:** Mariana L. Casalia, Tina Li, Harrison Ramsay, Pablo J. Ross, Mercedes F. Paredes, Scott C. Baraban

## Abstract

Interneurons contribute to the complexity of neural circuits and maintenance of normal brain function. Rodent interneurons originate in embryonic ganglionic eminences, but developmental origins in other species are less understood. Here, we show that transcription factor expression patterns in porcine embryonic subpallium are similar to rodents, delineating a distinct medial ganglionic eminence (MGE) progenitor domain. On the basis of Nkx2.1, Lhx6 and Dlx2 expression, *in vitro* differentiation into neurons expressing GABA and robust migratory capacity in explant assays, we propose that cortical and hippocampal interneurons originate from a porcine MGE region. Following xenotransplantation into adult male and female rat hippocampus, we further demonstrate that porcine MGE progenitors, like those from rodents, migrate and differentiate into morphologically distinct interneurons expressing GABA. Our findings reveal that basic rules for interneuron development are conserved across species, and that porcine embryonic MGE progenitors could serve as a valuable source for interneuron-based xenotransplantation therapies.

**Significance Statement:** Here we demonstrate that porcine MGE, like rodents, exhibit a distinct transcriptional and pallial interneuron-specific antibody profile, *in vitro* migratory capacity and are amenable to xenotransplantation. This is the first comprehensive examination of embryonic pallial interneuron origins in the pig, and because a rich neurodevelopmental literature on embryonic mouse MGE exists (with some additional characterizations in other species like monkey and human) our work allows direct neurodevelopmental comparisons with this literature.

## Introduction

Excitatory glutamatergic neurons and inhibitory GABAergic interneurons represent the two primary neuronal populations in mammalian brain. Cortical and hippocampal (pallial) inhibitory interneurons are a diverse subset of the total neuronal population, mediate many critical brain functions, and arise from embryonic subpallium regions (Fishell, 2007; Gelman et al., 2012; Kessaris et al., 2014). A wide spectrum of neurological pathologies is associated with loss, or dysfunction, of pallial interneurons including, but not limited to, epilepsy, autism spectrum disorder, Alzheimer’s disease and schizophrenia (Marin, 2012; Inan et al., 2013; Paterno et al., 2020). Studies over the past decade demonstrated that transplantation of pallial interneurons offer great promise for treatment of these disorders (Baraban et al., 2009; Alvarez-Buylla et al., 2000; Bráz et al., 2012; Anderson and Baraban, 2012; Hunt et al., 2013; Tong et al., 2014; Donegan et al., 2017; Juarez-Salinas et al., 2019; Zhu et al., 2019). Interneuron transplantation studies are largely based on harvesting progenitor cells from a medial ganglionic eminence (MGE) subregion. MGE progenitors give rise to cortical GABAergic interneurons subtypes expressing parvalbumin (PV)- and somatostatin (SOM) (Gelman et al., 2012; Hu et al., 2017; Pelkey et al., 2017) along with hippocampal neurogliaform and Ivy cells expressing nitric oxide synthase (nNOS) (Tricoire et al., 2010). These interneurons exhibit highly migratory capabilities and unique functions. To date, the majority of successful MGE progenitor cell transplantations showing efficacy in preclinical animal models harvest fresh murine embryonic tissue. Human “MGE-like” embryonic or induced pluripotent stem cell derived interneurons could offer an alternative, however, these neurons fail to (i) migrate extensively or (ii) differentiate to pallial interneuron subtypes seen with embryonic MGE progenitors, and (iii) exhibit protracted functional maturation (Nicholas et al., 2013; Liu et al., 2013; Maroof et al., 2014). Human stem cells also present a risk of tumorigenesis (Carpentino et al., 2008). Although embryonic allografted fetal human tissue in Parkinson’s disease patients ameliorated some symptoms of this disease in the 1990s (Björklund et al., 2003), practical and ethical problems using aborted human fetal tissue have since led to an extended exploration for an alternative source(s) of suitable fetal material for transplantation.

Most successfully applied to organs, xenotransplantation from pig-to-nonhuman primate (or even pig-to-human) raise the exciting possibility that embryonic porcine tissue - perhaps coupled with techniques to genetically modify a donor pig using transgenic or CRISPR/Cas9 technologies (Cowan et al., 2019; Hryhorowicz et al., 2020) - could be a viable source of transplantable pallial interneurons. These efforts are supported by data showing similarities at the gene and protein level between humans and pigs (Sjöstedt et al., 2020). It is well established that murine MGE can be defined by transient anatomical landmarks and expression of a unique set of transcription factors e.g., Nkx2.1, Dlx2 and Lhx6 (Corbin et al., 2003; Du et al., 2008; Flames et al., 2007; Gelman et al., 2012). Analysis in human and macaque (nonhuman Old World) monkey demonstrated a recapitulation of these murine MGE transcription factor expression patterns (Hansen et al., 2013; Ma et al., 2013). Tangential migration of pallial interneurons from MGE-to-cortex or -hippocampus is also believed to be highly conserved across species. Although some features of porcine ganglionic eminences are known, this information is limited to lateral ganglionic eminence (LGE) (Jacoby et al., 1999), a subpallial region that mainly generates olfactory bulb interneurons and lacks tangential migratory properties *in vitro* or following transplantation into postnatal brain.

Here we analyzed temporal and regional expression of MGE-specific transcription factors in fetal porcine brain at embryonic days 30 and 35 (E35). Cultured porcine progenitors express MGE-specific transcription factors, differentiate to GABAergic SOM^+^ interneurons and migrate extensively *in vitro*. Similar to human, embryonic pig ganglionic eminence organization showed distinct organization of cells into doublecortin-positive clusters, a pattern not seen in murine MGE. Xenotransplantation of E35 MGE progenitors into adult rat hippocampus resulted in migration across all hippocampal sub-fields and differentiation into GABAergic SOM^+^ interneurons up to 60 days after transplantation (DAT). Overall, patterns of transcription factor expression, differentiation into distinct interneuron subpopulations and tangential migration capacity all appear to be conserved from rodents to pig MGE during evolution.

## Materials and Methods

### Animals

Donor porcine (*Sus scrofa domesticus*) embryos were procured from Department of Animal Science, University of California, Davis (English Large white) or National Swine Resource and Research Center (NSRRC:0016 GFP *NT92*; Whitworth et al., 2009), University of Missouri. Adult male Sprague Dawley rats (*Rattus norvegicus*) were purchased from Charles River Laboratories (strain code 400) and housed under a standard 12 hr light/dark cycle with food and water provided *ad libitum*. A total of 133 rats were used for xenotransplantation studies (79 female; 54 male); 75 rats received intraperitoneal (i.p.) injection of an immunosuppression cocktail containing methylprednisolone acetate (2 mg/kg), cyclosporine (20 mg/kg) and azathioprine (5 mg/kg) 3 times per week (Table 1). All protocols and procedures were approved by the Institutional Animal Care and Use Committee (IACUC protocol #AN181524-02) and adhered to University of California, San Francisco (UCSF; San Francisco, CA) Laboratory Animal Resource Center and United States Public Health policy on Humane Care and Use of Laboratory Animals guidelines.

**Table 1:** Porcine MGE donor cell manipulations.

| donor cells | post hoc histology |  | total (n) | transplant survival rate (%) |
| --- | --- | --- | --- | --- |
|  | with cells | no cells |  |  |
| pMGE wt + lenti CMV GFP / immunosup | 10 | 15 | 25 | 40 |
| pMGE wt + lenti CMV GFP / NO immunosup | 1 | 19 | 20 | 5 |
| pMGE GFP / immunosup | 19 | 31 | 50 | 38 |

\*immunosuppression cocktail: methylprednisolone acetate (2 mg/kg), cyclosporine (20 mg/kg) and azathioprine (5 mg/kg)

### Tissue collection and processing

Individual embryonic day 30 (E30) or E35 pig embryos were maintained on ice in 50 ml Falcon conical centrifuge tubes (Fisher Scientific, catalog #14-432-22) with DMEM/F12 media. Sterile techniques were used during isolation and microdissection. Cell viability on freshly dissected brain tissue was assessed using Trypan Blue (Sigma); only tissue with ≥ 90% viability were processed for cryopreservation.

### Immunohistochemistry

Embryonic pig heads were removed under a stereomicroscope (Leica MZ12(5)) and placed individually in a 50 ml Falcon tube containing 30 ml of 4% paraformaldehyde (PFA; wt./vol) in phosphate buffered saline (PBS) at 4°C for 48 hr. Rats were deeply anesthetized and perfused with 300 ml of cold 4% paraformaldehyde solution. Brains were removed and post fixed ON with 4% PFA. Tissue was sequentially dehydrated and cryoprotected in 10%, 20% and 30% sucrose (wt./vol) in PBS at 4°C until sinking. Tissue was embedded in O.C.T. compound (Tissue-Tek #4583) and frozen on a dry ice/ethanol slush and stored at −80°C for serial sectioning on a Leica CM1900 cryostat. Thin tissue sections (30 μm for pig tissue; 25 μm for rat tissue) were collected onto glass slides (SuperFrost Plus, catalog #12-550-15) or free floating in cryopreservation solution (3 vol. glycerol, 3 vol. ethylene glycol, 4 vol 1x 0.1% Sodium Azide in PBS) and stored at −20°C. Tissue on SuperFrost slides were stained using TSA amplification kit (Perkin Elmer; # NEL701A001KT, # NEL704A001KT and # NEL705A001KT). Briefly, cryosections were left at ON at RT, baked for 15 min at 60°C, fixed for 15 min with 4% PFA. Antigen retrieval was performed using nearly boiling 10 mM sodium citrate solution (5.95-6.05 pH range). Sections were allowed to cool down to RT and tissue was delineated with pap-pen (Abcam, # ab2601). Free floating sections were left at RT for 2 hr prior 3 × 15 min washes with PBS to remove cryopreservation solution. Tissue was double stained with a primary antibody and detected with Alexa secondary antibodies (Table 2). After testing different commercially available antibodies, we failed to identify one that could reliably distinguish pig PV^+^, calbindin^+^, calretinin^+^, reelin^+^ or VIP^+^ interneurons. Nuclei staining was done for 5 min as a final step (Sigma; H33342). Briefly, sections were incubated with 0.3% triton-x 100 in PBS for 60 min at RT, incubated in blocking solution (10% Donkey serum, 1% BSA, 0,1% Triton-X 100 in PBS) for 60 min and incubated ON in primary antibody at 4° C. The next day sections were washed 3 x 10 min in 0.05% Triton-X 100 in PBS, incubated for 120 min at RT in secondary antibodies. Sections were mounted onto charged slides (Fisher Scientific, Superfrost Plus) using Fluoromount-G solution (Southern Biotech #0100-01).

**Table 2:** Antibodies.

| primary antibodies | host specie | company | catalog # | dilution |
| --- | --- | --- | --- | --- |
| CoupTFII | mouse | Perseus Proteomics | PP-H7147-00 | 1/100 |
| DBX1 | rabbit | abcam | ab156283 | 1/500 |
| DCX | mouse | Santa Cruz Biotechnology | sc-271390 | 1/100 |
| Dlx2 | mouse | Santa Cruz Biotechnology | sc-393879 | 1/500 |
| GABA | rabbit | sigma | a2052 | 1/500 |
| GFAP | mouse | EnCore | MCA-5C10 | 1/1000 |
| GFAP | rabbit | Dako | Z0334 | 1/500 |
| KI67 | mouse | BD pharmingen | 556003 | 1/500 |
| KI67 | rabbit | proteintech | 27309-1-AP | 1/1000 |
| LHX6 | rabbit | Santa Cruz Biotechnology | sc-98607 | 1/100 |
| LHX6 | mouse | Santa Cruz Biotechnology | sc-271433 | 1/500 |
| MAP2 | mouse | EnCore | MCA-4h5 | 1/1000 |
| Nkx2.1 (TTF1) | mouse | Progen | 16108 | 1/20 |
| Olig2 | rabbit | EMD Millipore | AB9610 | 1/500 |
| Pax6 | mouse | abcam | ab78545 | 1/500 |
| PSA-NCAM (IgM isotype) | mouse | millipore | MAB5324 | 1/500 |
| PV | rabbit | swant | PV27 | n/a |
| PV | rabbit | abcam | ab11427 | n/a |
| PV | chicken | EnCore | CPCA-Pvalb | n/a |
| Somatostatin | mouse | Santa Cruz Biotechnology | sc55565 | 1/500 |
| Somatostatin | goat | Santa Cruz Biotechnology | sc-7819 | 1/200 |
| Synaptophysin | mouse | Santa Cruz Biotechnology | sc17750 | 1/100 |
| Tau | chicken | EnCore | CPCA-Tau | 1/500 |
| Tuj (BIII tubulin) | rabbit | covance | MRB-435P | 1/1000 |
| VGAT | mouse | Santa Cruz Biotechnology | sc 393373 | 1/100 |
| VIP | Rabbit | abcam | AB22736 | n/a |
| nNOS | rabbit | millipore | AB5380 | n/a |
| Calbindin | rabbit | Chemicon | AB1778 | n/a |
| Calretinin | mouse | Chemicon | MAB1568 | n/a |
| Calretinin | rabbit | Millipore | AB5054 | n/a |
| Calretinin | Mouse | Swant | 7697 | n/a |
| reelin | mouse | millipore | MAB5364 | n/a |
| Reelin | Mouse | Calbiochem | 553730 | n/a |
| Neuropeptide Y | Rabbit | Abcam | ab30914 | n/a |
| **amplification system for E35 stains** |  |  |  |  |
| TSA Biotin System |  | PerkinElmer | NEL700A001KT |  |

### Image processing and cell quantification

Images were acquired from fluorescently labeled sections using Keyence B7-X710, Leica TCS SP5 or Nikon Eclipse Ni-E confocal microscopes. For whole-brain visualization, images were acquired using a 10x objective in Tilescan mode on a Keyence microscope with Z-stack to focus automatic stitching of arrayed tiles. Live imaging movies were acquired on a Leica TCS SP5 using 10x objective, 1.5x optical zoom in time intervals of 20 min. All labeled cells were counted using FIJI (http://imagej.net) by an investigator blind to the status of the experiment. Gray level intensity was set using thresholded binary images in FIJI Watershed.

### Experimental Design and Statistical Analysis

#### Cell culture

MGE was dissociated into single cells using neuronal isolation enzyme mix, as described previously^45^. A 50 μl drop of 10% FBS in NIM (DMEM Ff12, NEAA 1:100,N2 1x, Heparin 1:1000, B27 1x, EGF, FGF) containing 5,000 cells/cm^2^ was placed onto a polyornithine/laminin coated coverslip in 24-well plate for 4 hr to allow initial attachment. An additional 100 μl of 10% FBS in NIM was added for overnight (ON) incubation and further attachment of cells to the coverslip. Next morning, all media containing FBS was removed and fresh NIM with no FBS with factors (BDNF, GDNF, IGF and cAMP 1:5000 vol/vol) was added to start the differentiation process. Cells were allowed to differentiate for 14 days with media change every 48 hr. Factors were freshly added to the media. After cells reached desired differentiation stage, they were fixed in 4% PFA for 5 min at room temperature (RT) and stored in 0.1% sodium azide (wt/vol) in PBS for histology.

#### Xenotransplantation

MGE was identified by anatomical landmarks and freshly dissected, as described (Hunt et al., 2013; Casalia et al., 2017) (Fig. 9). Tissue was then cryopreserved in 10% DMSO in DMEM/F12 using 1.5 ml cryotubes and stored in a liquid nitrogen tank between −210°C and −196°C. On the day of xenotransplantation, cryotubes were quickly thawed under running warm water with gentle shaking. Neuronal Isolation Enzyme mix (Pierce #88285E) was used for cell dissociation (30 min at 37°C). Cell viability on freshly thawed cells was assessed using Trypan Blue; only MGE progenitor cells with ≥ 80% viability were processed for xenotransplantation. Rats were anesthetized with 1.5% to 2.5% isoflurane in oxygen (vol./vol.) and secured in a stereotaxic frame (Kopf model #900). In a series of pilot studies, several needle sizes and cell volumes were tested (Fig. 10). A protocol using single cell suspension of 50,000 cells in 0.5 μl of DMEM/F12 loaded into a 32G blunt Hamilton syringe was chosen. Target coordinates for stratum radiatum of hippocampal CA3 subfield were initially verified in a series of pilot dye injections (n = 3) at the following stereotaxic coordinates relative to bregma: anterior-posterior (AP) −3.4; medial-lateral (ML) ± 2.7; dorsal-ventral (DV) −3.4. Cell deposits were made using an automatic microinjection pump (WPI Microinjection System UMP3) at a rate of 5 nl/sec. The needle remained in place for 60 sec and then slowly withdrawn. Following xenotransplantation, the incision was closed with Vetbond (3M #1469SB) and rats were placed under a heat lamp until fully recovered. A successful porcine MGE xenotransplant was defined *post hoc* using histological techniques. Namely, the presence of at least 40 GFP^+^ cells in a minimum of 2 sections of the hippocampus (25 μm thick sections, spaced 300 μm apart).

#### Explant assay

MGE was dissected and further sub-dissected in 4 sections (anterior-dorsal (AD)/ anterior-ventral (AV)/ posterior-dorsal (PD), posterior-ventral (PV), see schematic figure 6). Each sub-section was submerged in a Matrigel drop (Corning, # 356237) containing 50% of cell culture media (Neurobasal, 1x amphotericin, 1x penicillin-streptomycin, 1x glutaMAX) and placed onto a Matrigel coated coverslip of a 24-well plate at 37° C under hypoxic conditions (5% CO_2_, 8% O_2_ balanced N_2_). Explants were allowed to attach ON with addition of 10% FBS to cell culture media. Next morning media containing FBS was removed and replaced with serum free media. Explants were kept *in vitro* for 6 days with media change every 48 hr. At the end of the process each coverslip was carefully washed 3 times with PBS and fixed for 15 min in 4% PFA in PBS solution for histological analysis.

#### Slice culture

E35 brains were removed out of the head in cold DMEM F12 media. Tissue was embedded in 3.5% low melting point agarose in 310 mOsm artificial cerebrospinal fluid (ACSF; 125 mM NaCl, 2.5 mM KCl,1 mM MgCl_2_, 2 mM CaCl_2_, 1.25 mM NaH_2_PO4, 25 mM glucose, bubbled with 95% O_2_ / 5% CO_2_) and sliced using a Leica VT1200S vibratome at 300 μm thickness. Sections were allowed to stabilize for 20 min in bubbling ACSF on ice before transferred and suspended on 0.4 μm Millicell-CM slice culture inserts (Millipore). Sections were cultured for 6 days at 37°C in 5% CO_2_, 8% O_2_ and balanced N_2_. For time-lapse imaging, an adenovirus (AV-CMV-GFP, 1 × 1010; Vector Biolabs) at a dilution of 1:50–1:500 was applied to the slices, which were then cultured at 37°C, 5% CO_2_, 8% O_2_. For time-lapse imaging, cultures were then transferred to an inverted Leica TCS SP5 confocal microscope with an on-stage incubator streaming 5% CO_2_, 5% O_2_, and balanced N_2_ into the chamber. Slices were imaged using a 10× air objective at 20-minute intervals for 72 hours with repositioning of the z-stacks every 6-8 hr.

#### Statistics

Data is expressed as mean ± s.e.m. (standard error of the mean). Statistical significance was assessed using one- or two-way ANOVA with Kruskal-Wallis’s multiple comparisons. *P*-values less than < 0.05 were considered significant.

## Results

### Temporal and regional identification of porcine embryonic MGE

Across several species (Lavdas et al., 1999; Metin et al., 2007; Tanaka et al., 2011; Gelman et al., 2012; Hansen et al., 2013; Ma et al., 2013; Hu et al., 2017), ganglionic eminences (GE) consisting of extensive proliferative cell masses are visible as distinct elevations adjacent to, and protruding into, lateral ventricular walls. Identification of embryonic lateral and medial GE subregions is guided by transient developmental appearance of anatomical landmarks (Brazel et al., 2003; Flames et al., 2007). In serial sections, an expansion of porcine embryonic forebrain occurs between E30-E35 along with formation of a clear morphological boundary e.g., a sulcus in the ventricular surface separating LGE and MGE (Fig. 1). GE subregions can also be identified based on temporal transcription factor expression patterns. **Nkx2.1**, a homeobox transcription factor gene, is required for MGE specification and highly expressed in early MGE ventricular zones (Corbin et al., 2003; Du et al., 2008; Sandberg et al., 2016). At E30 and E35, we observed prominent anterior-to-posterior Nkx2.1 expression coinciding with appearance of a sulcus and anatomical designation of an MGE subregion (Fig. 1a, *arrows*); absence of expression in LGE and CGE was also noted (Fig. 1b). **Gsh2** (also known as Gsx2), another homeobox transcription factor gene, is critical for dorsal-ventral patterning of telencephalon and diffusely expressed throughout GE domains (Flames et al., 2007; Du et al., 2008). Coronal sections at E30 show diffuse expression of Gsh2 throughout porcine embryonic GE while evolving to more restricted expression patterns corresponding to CGE and LGE ventricular proliferative zone regions at E35 (Figs. 1a-b, *white arrows*). **Olig2**, a basic helix-loop-helix transcription factor gene, is expressed in progenitor cells of ventral GE with highest levels in Nkx2.1-expressing MGE subregions (Miyoshi et al., 2007). At E35, Olig2 expression is prominent in proliferative zones of porcine MGE with only sparse expression in CGE or LGE (Fig. 1b). **Dlx** homeobox transcription factors (Dlx1/2 and Dlx5/6), widely expressed throughout subpallium, are broadly required for GE progenitors to migrate and differentiate into GABAergic interneurons (Flames et al., 2007; Du et al., 2008; Laclef and Metin, 2018). Porcine Dlx2 expression at E35 is prominent in MGE (and CGE) though somewhat less dense along ventricular proliferative zones. LGE expression is more diffuse presumably because interneuron progenitors begin migration toward cortical areas at this stage of embryonic development. Optical density measurements on coronal sections confirm high levels of Dlx2 and Olig2 expression in all three ventricular zones of porcine MGE at E30 and E35 (Fig. 1c). **Lhx6**, a LIM homeodomain transcription factor, activated by Nkx2.1 also plays a critical role in specification of MGE-derived interneuron subtypes (Grigoriou et al., 1998; Du et al., 2008). Lhx6, expressed in MGE subventricular and submantle zones, is required for differentiation of PV+ and SOM+ interneurons^36,37^. At E30, we observed significant Lhx6 expression as a column of migrating cells bordering ventricular zones, contiguous with basal MGE and beginning to populate piriform cortex (pCx) ventrally and LGE laterally (Fig. 2a, *coronal* and *sagittal*). Dlx2 shows a similar co-expression pattern but overall density is less in piriform cortex and scattered expression in ventricular zones was noted. At E35, Lhx6 expression in presumably migratory cells outside the MGE ventricular zone is more prominent ventrally (Fig. 2a, *coronal*) along with scattered expression into intermediate zones of LGE and cortex (Fig. 2a, *coronal* and *sagittal*; Fig. 3b). Dlx2 expression at E35 is prominent in MGE and LGE subregions. Nkx2.1 co-expression with Lhx6 or Dlx2 can be seen in most cells within the MGE ventricular proliferative zone at E35 (Fig. 2b1, *coronal*; Fig. 3a) but only a few cells in more intermediate zones (Fig. 2b2, *sagittal*). Serial coronal sections along the anterior-to-posterior axis (Fig. 2c) reveal distinct Lhx6 expression in presumed tangential telencephalon-to-cortex, telencephalon-to-striatum, telencephalon-to-olfactory bulb migratory streams, as previously described in mice (Marin and Rubenstein, 2001; Alifragis et al., 2004; Liodis et al., 2007); Lhx6 co-expression with Dlx2 was also noted in MGE (Fig. 3a). In medial coronal sections at E35, MGE (marked by intense Nkx2.1 and Lhx6 expression) co-expressed Dlx2 and Olig2 more densely than LGE (Figs. 3a-b). Expression of CouptfII (an orphan nuclear receptor enriched in CGE)(Kanatani et al., 2008) was expressed in dorsal MGE, as expected (Cai et al., 2013); Gsh2 (enriched in LGE/CGE)(Du et al., 2008; Laclef and Metin, 2018) was not prominent in MGE (Fig. 3b). Non-specific ventral forebrain progenitor markers (Ki67, Olig2 and DCX) label porcine MGE at E35 with dense co-expression of Olig2 and Ki67 noted (Fig. 3b).

**Figure 1:**
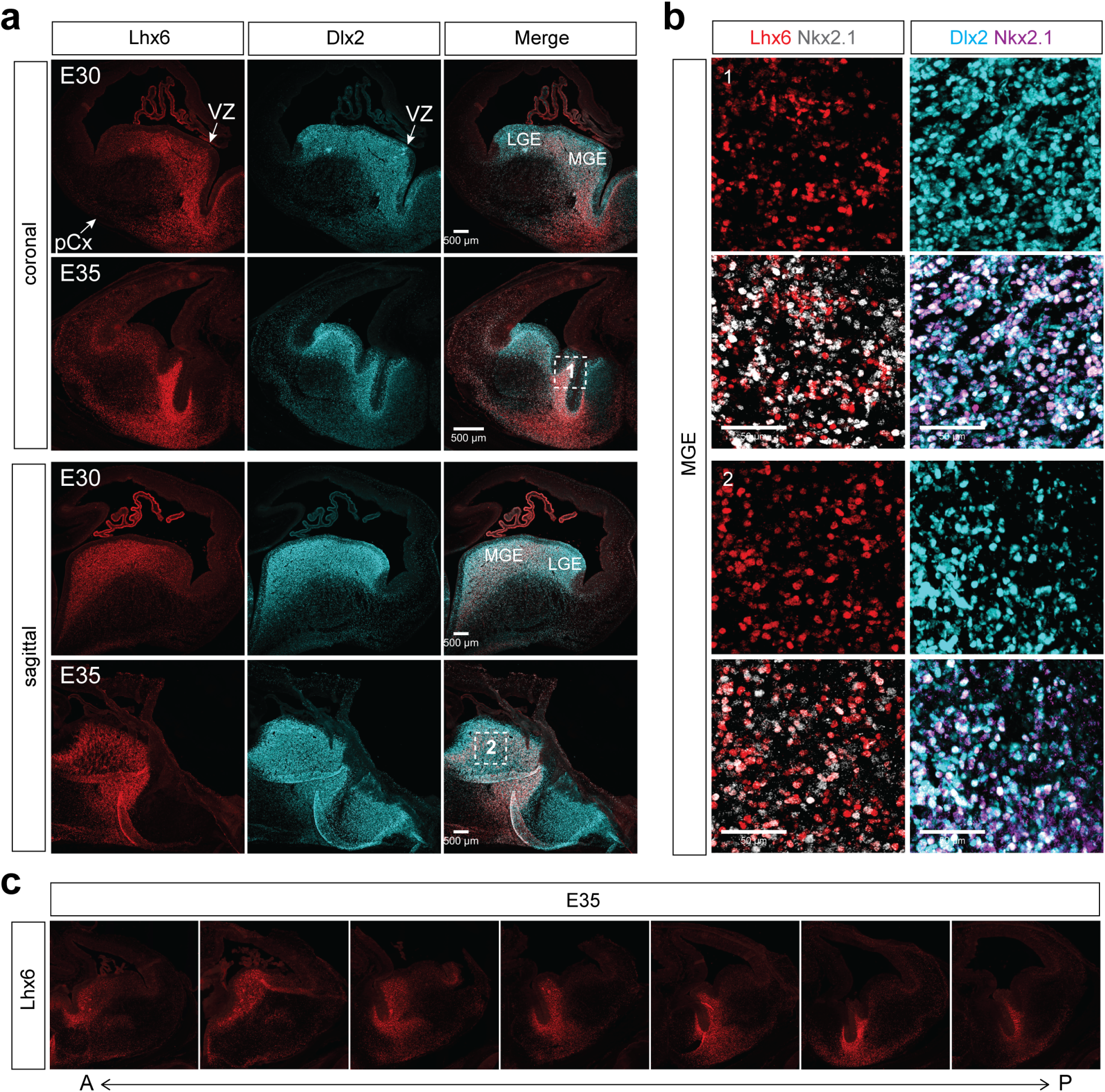
Developmental expansion of ganglionic eminences in the pig embryonic brain. (**a**) Serial coronal sections of E30 and E35 pig brain along the anterior (A) to posterior (P) axis. Sections labeled with antibodies recognizing regional transcription factors that specify ganglionic eminences, Nkx2.1 (*red*) and Gsh2 (*purple*). DAPI cell stain shown in *blue*. Arrows distinguish an anatomically defined MGE subregion; sulcus establishing a boundary between LGE and MGE became apparent at E35 (*yellow arrows*) and Gsh2 expression in LGE and CGE subventricular zone (*white arrows*) (E30, n = 9; E35, n = 9). (**b**) Regional (Nkx2.1, Gsh2) and temporal (Olig2, Dlx2) transcription factor expression at E35 for MGE, LGE, and CGE subregions (n = 9). Note distinct MGE-specific Nkx2.1 expression. Scale bar = 100 μm (**c**) Grouped summary data plots showing intermediate progenitor (Dlx2) and undifferentiated radial glia (Olig2) expression at E30 and E35 (n = 3). Schematic (inset) shows where ventricular zone (VZ), subventricular zone 1 (SVZ1) and 2 (SVZ2) optical density (O.D.) measurements were made.

**Figure 2:**
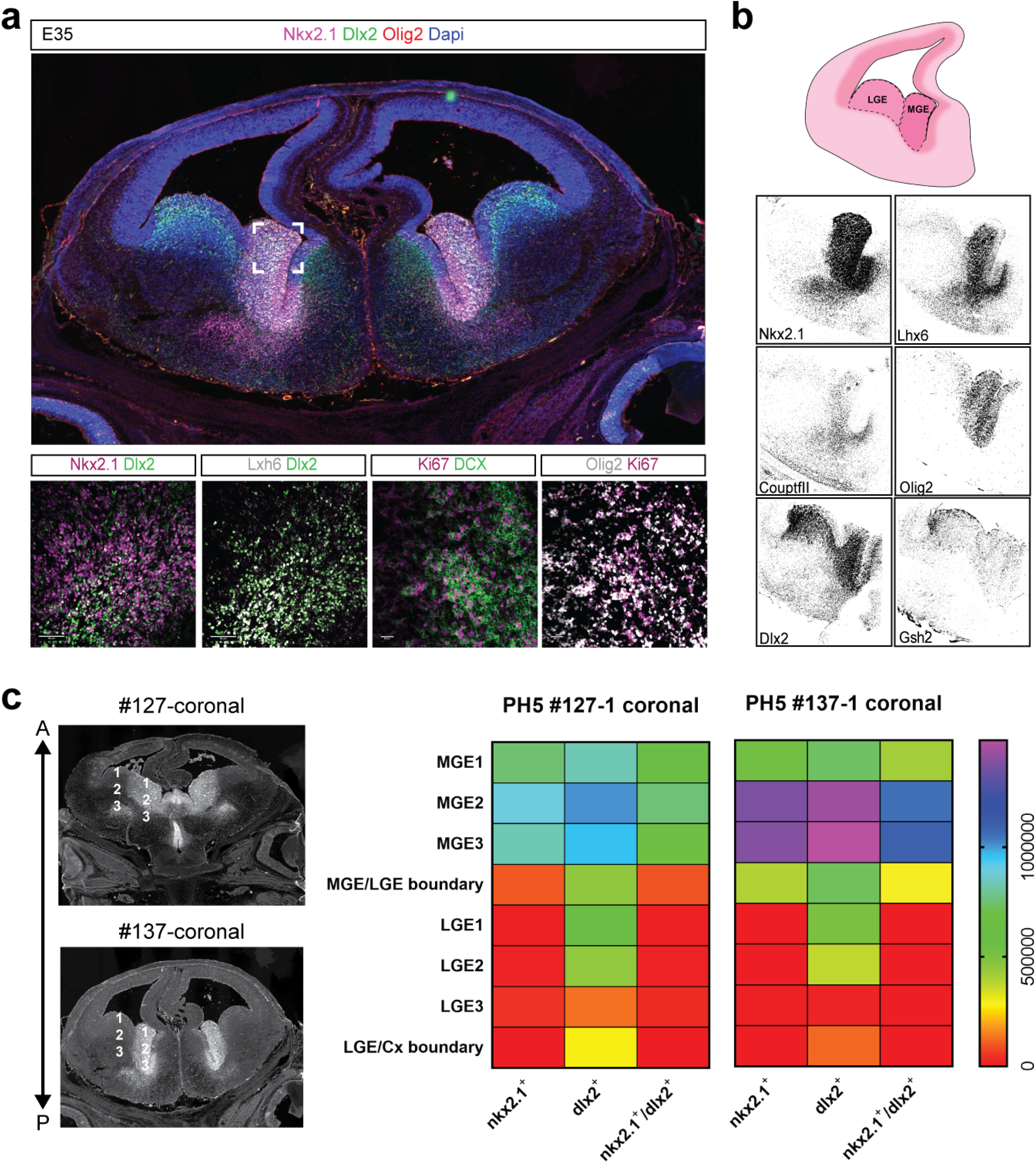
MGE-specific transcription factor expression the developing porcine ventral forebrain. (**a**) Representative coronal and sagittal sections containing the medial GE region are shown at E30 and E35. Dlx2-expressing intermediate and Lhx6-expression progenitors exit the ventricular zone (VZ) to begin a tangential migration toward cortical destinations (n = 9). Scale bar = 500 μm. (**b**) Higher magnification of inset 1 and 2 of panel (a) showing MGE expression of Nkx2.1 expression. Note high co-expression of Nxk2.1 and Lhx6 (left panels) or Dlx2 (right panels), in coronal and sagittal MGE sections. Scale bar = 50 μm (**c**) Serial coronal sections of E35 brain delineating Lhx6 expression corresponding to MGE progenitor migratory routes along the anterior-to-posterior axis (n = 9). Abbreviations: ventricular zone (VZ), medial ganglionic eminence (MGE), lateral ganglionic eminence (LGE) and piriform cortex (pCX).

**Figure 3:**
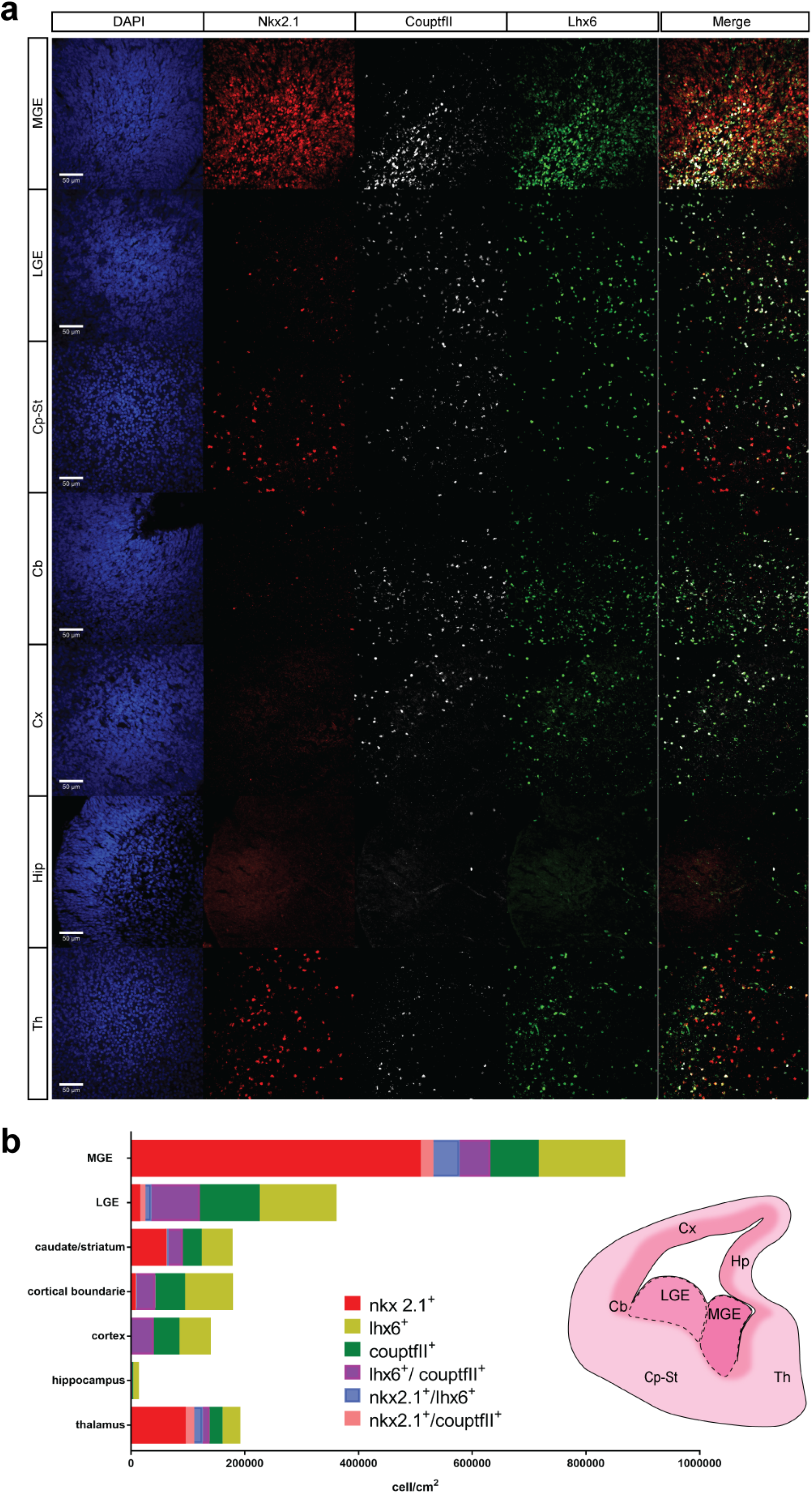
Demarcation of an embryonic MGE subregion in the developing pig brain. (**a**) Co-expression of Nxk2.1, Dlx2 and Olig2 in a representative coronal brain section at E35 (top panel). Higher resolution images shown in lower panels (box) highlight co-expression patterns for intermediate progenitors (Nkx2.1 and Dlx2), MGE-specific transcription factors (Lhx6 and Dlx2), proliferating and young neurons (Ki67 and DCX) or MGE-derived oligodendrocytes and proliferating cells (Olig2 and Ki67) (n = 9). (**b**) Schematic showing localization of MGE and LGE subregions on an equivalent section corresponding to the gray scale images (top panel). Transcription factor markers (Lhx6 and Dlx2) label the porcine MGE (marked by Nkx2.1). Olig2 expression is also prominent in MGE. CouptfII expression overlaps with migratory cells exiting the MGE. Gsh2 expression in VZ regions are most prominently in LGE (n = 18). (**c**) Representative coronal sections (#127 & #137) along the Anterior-to-Posterior at 300 μm are shown at E35 brain. Nxk2.1^+^, Dlx2^+^ and Nkx2.1:Dlx2^+^ expressing cells were counted. MGE and LGE subregions, designated numbers 1-3, were used for cell density measurements (in cells/cm^2^). Heatmaps of these GE region cell density measurements are shown at left. Heatmap color scale = 0 – 150,000 cells/cm^2^ (n = 18).

Next, we performed a compartmentalized quantitative analysis of Nxk2.1 and Dlx2 expressing cells using coronal sections representative of anterior or posterior levels of E35 porcine MGE. MGE and LGE were divided into 3 subregions starting at the ventricular surface (MGE1) as well as boundary regions for MGE/LGE or LGE/Cortex (Fig. 3c). In the most anterior section (Fig. 3c, PH5 #127-1), Nkx2.1 cell density is highest in MGE regions 1 and 2 corresponding to proliferative ventricular zones, and extending into subregion 3 of the most posterior section (Fig. 3c, PH5 #137-1). At a posterior level, Nkx2.1 cell density drops off at the MGE/LGE boundary and is absent from LGE and LGE/Cortex boundaries. Dlx2 cell density is high throughout all MGE proliferative zones and subregions, MGE/LGE boundary and into LGE subregions 1 and 2 (Fig. 3c). As expected (Flames et al., 2007; Long et al., 2009), co-expression of cells expressing Nkx2.1 and Dlx2 cells was restricted to MGE at both levels.

It is well established that MGE produces GABAergic neurons that tangentially migrate to hippocampus, cortex, striatum and thalamus (Marin and Rubenstein, 2001; Metin et al., 2007; Laclef and Metin, 2018). Next, we used down-regulation of Nkx2.1 expression in migratory cells to delineate MGE regions coupled with Lhx6 expression to track migratory cells and the transition between germinal zone and migratory routes at E35 (Fig. 4). Large populations of Nkx2.1^+^ progenitor cells were observed in MGE with additional Nkx2.1^+^ cells migrating into caudate-striatum and thalamus. Nkx2.1^+^ cell density greatly decreased in LGE consistent with down-regulation of this transcription factor after leaving MGE (Marin and Rubenstein, 2001); Nkx2.1^+^ cells were not observed in cortex or hippocampus. A significant population of Lhx6^+^ cells can be seen in dorsal MGE along with CouptfII^+^ cells (Fig. 4). Outside MGE, similar patterns of Lhx6^+^ and CouptfII^+^ cells can be seen in caudate-striatum, thalamus, and cortex suggesting a corridor for tangential migration. Distinct non-overlapping populations of Nkx2.1^+^ and Lhx6^+^/CouptfII^+^ cells are present in caudate-putamen and thalamus. Taken together, these observations support a conclusion that temporal, morphological and molecular characteristics of embryonic MGE are conserved in the pig.

**Figure 4:**
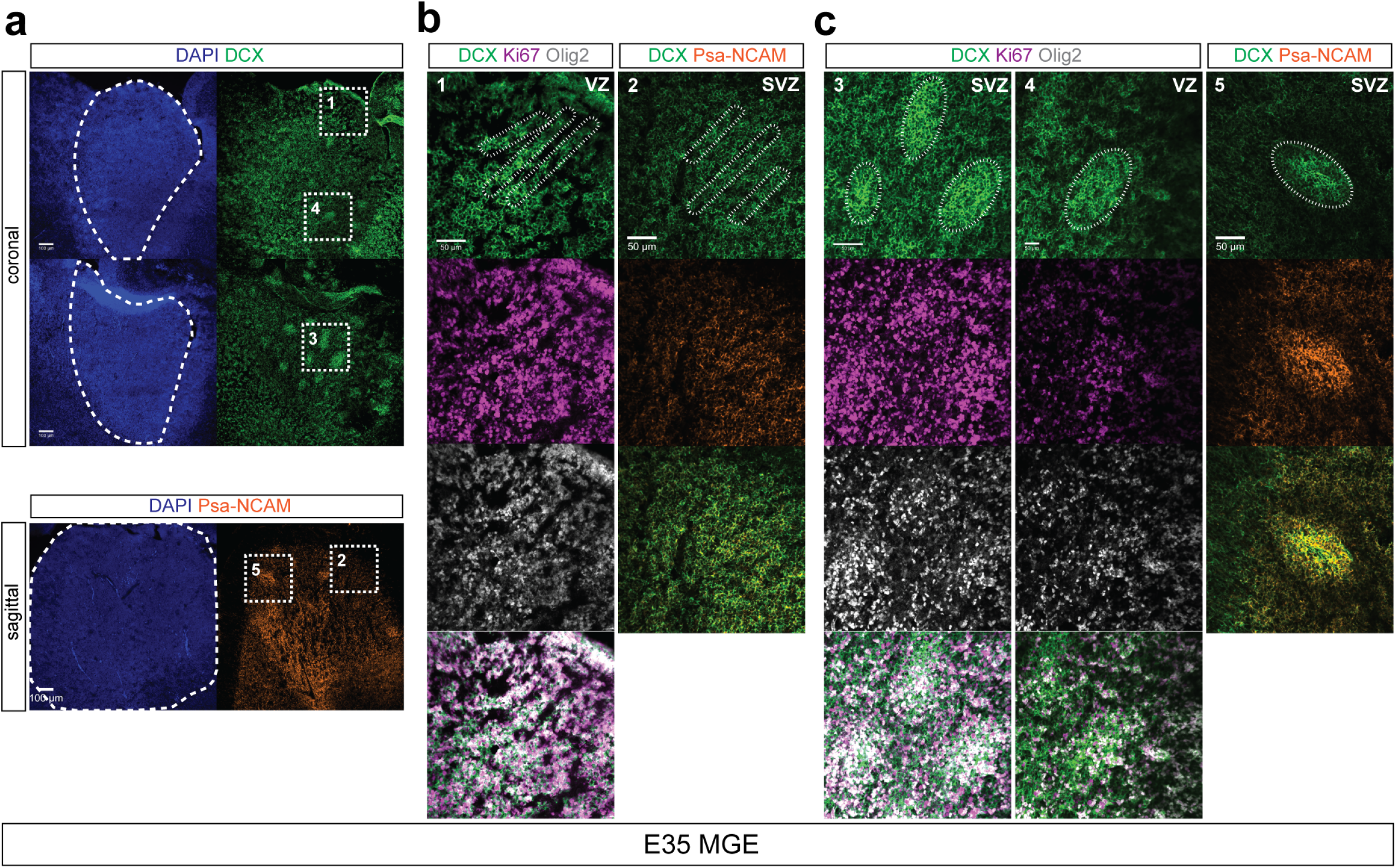
MGE-derived progenitors cell distribution in porcine embryonic subpallium. (**a**) Representative Nkx2.1, CouptfII and Lhx6 expression patterns in porcine at E35 are shown. DAPI stain and merged images are also shown (n = 9) (scale bar = 100 μm). (**b**) Grouped summary data plots showing transcription factor (Nxk2.1, Lhx6 and CouptfII) cell densities (cell/cm^2^) for embryonic subregions shown in schematic (at right). Co-expression quantification is also shown for Lhx6:CouptfII^+^, Nkx2.1:Lhx6^+^ and Nkx2.1:CouptfII^+^ cells in these subregions (n = 7). Abbreviations: MGE, medial ganglionic eminence; LGE, lateral ganglionic eminence; Cp-St, caudate putamen-Striatum; Cb, cortical boundary; Cx, cortex; Hip, hippocampus; and Th, Thalamus.

At E35, we also noted a unique organization of DCX^+^ cells originating close to the ventricle in VZ or SVZ regions of porcine MGE (Figs. 5a-b, inset #1). Organization of MGE progenitor cells into clusters was described for human and primate (but not murine) embryonic brains (Hansen et al., 2013; Ma et al., 2013). Some co-expression with a marker of young proliferative Ki67^+^ cells was observed in these DCX^+^ cluster-like formations. These cells extended transversally towards outer ventricular areas with co-expression of Psa-NCAM, a neural cell adhesion molecule regulating maturation of GABAergic interneurons^42^ (Figs. 5a-b, inset #2). More distinct oval-shaped clusters were seen in outer VZ and SVZ further from the ventricular surface (Figs. 5a, c, insets #3-5). Oval-shaped clusters were densely populated by a core of DCX^+^/Ki67^+^proliferative young neurons and clear co-expression of Olig2 and Psa-NCAM.

**Figure 5:**
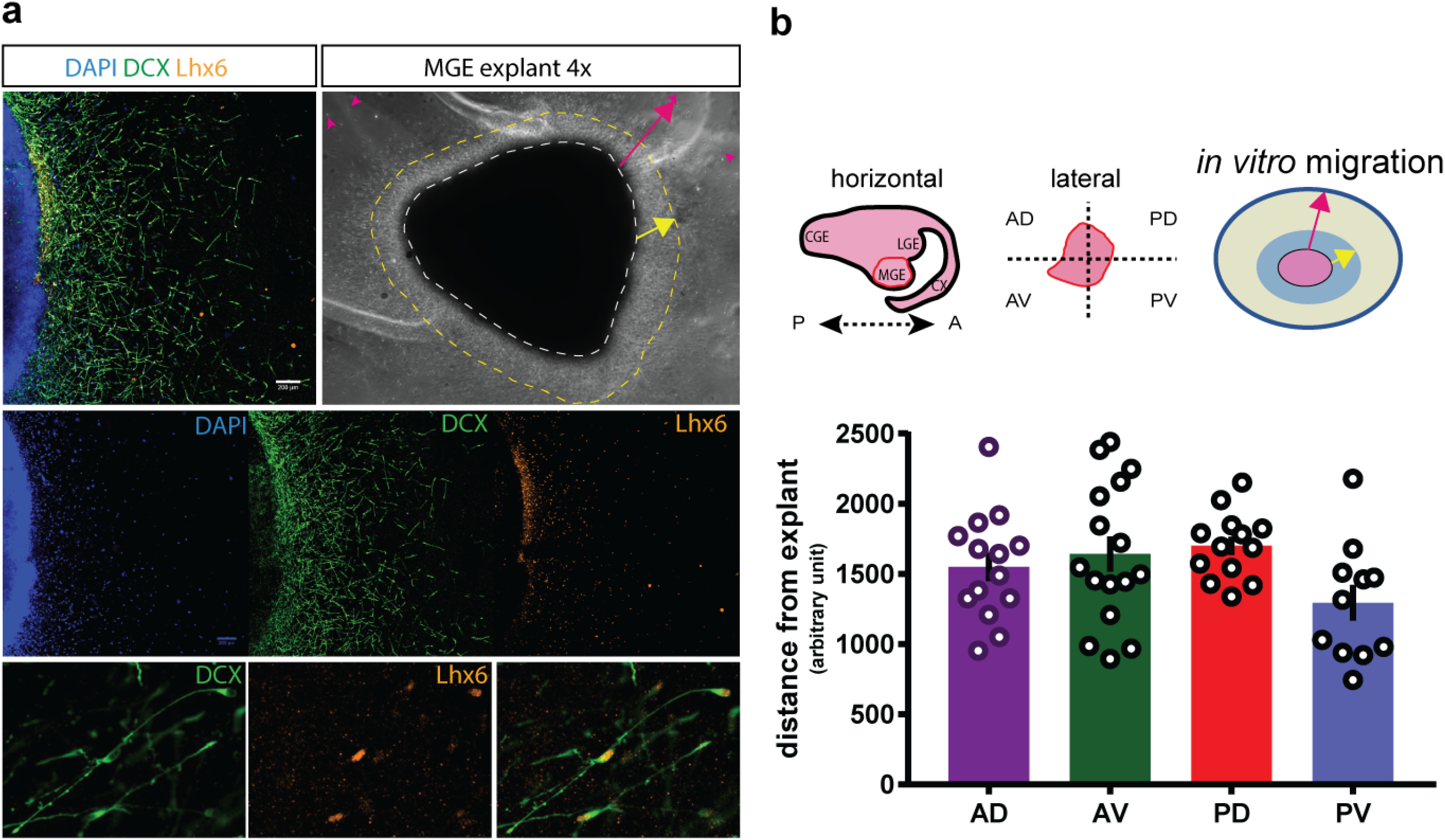
Organization of DCX^+^ clusters in embryonic porcine MGE at E35. (**a**) Representative coronal and sagittal sections showing DCX clustering in the MGE (designated by white boundary). Note DCX^+^ or Psa-NCAM^+^ cell organization in panels at left. (**b**) Magnified images of boxed areas in (a) show organization of cells co-expressing Ki67 and Olig2 (#1) or DCX and Psa-NCAM (#2). (**c**) Magnified images of boxed areas in show clusters of cells co-expressing DCX, Ki67 and Olig2 (#3-4, subventricular and ventricular zones, respectively) or DXC and Psa-NCAM (#5). Scale bar = 50 μm. (n = 18)

### *In vitro* migration and characterization of porcine MGE

Embryonic MGE progenitor cells exhibit chain migration when cultured in a three-dimensional extracellular matrix gel (Matrigel)(Wichterle et al., 1997, 2003). Using the same explant protocol, E35 porcine MGE sections were placed in Matrigel drops and kept under hypoxic conditions for 6 days *in vitro* (6 DIV). Extensive migration from MGE explants, forming an outer ring of cells emerging from the core, was seen as early as 3 DIV, and grew more extensive by 6 DIV (Fig. 6a, *yellow arrow*). Outside this band cells were loosely arranged forming a meshwork of migrating cell chains emanating from the explant core in all directions (Fig. 6a, *magenta arrow*) and reaching distant areas of the plate (Fig. 6a, *magenta arrowhead*). Cells migrating outward from explants co-expressed: (i) DCX, a microtubule-associated protein expressed by neuronal progenitor cells and immature neurons and (ii) Lhx6, an MGE-specific transcription factor (Fig. 6a, middle and lower panels). DCX^+^ cells had an elongated morphology resembling those described for migrating mouse MGE cells *in* vitro (Wichterle et al., 1997). We also observed robust individual cell migration in time-lapse movies obtained from E35 porcine MGE tissue slices cultured *in vitro* for 48 hours (Supplemental Movie). To quantify migration capacity, MGE explants were sub-divided into Anterior or Posterior and Dorsal or Ventral regions (AD/PD/AV/PV) and individual cell distance from explants was measured (Fig. 6b, schematic). As expected (Wichterle et al., 1997, 2003), migration was robust in every sub-region (Fig. 6b).

**Figure 6:**
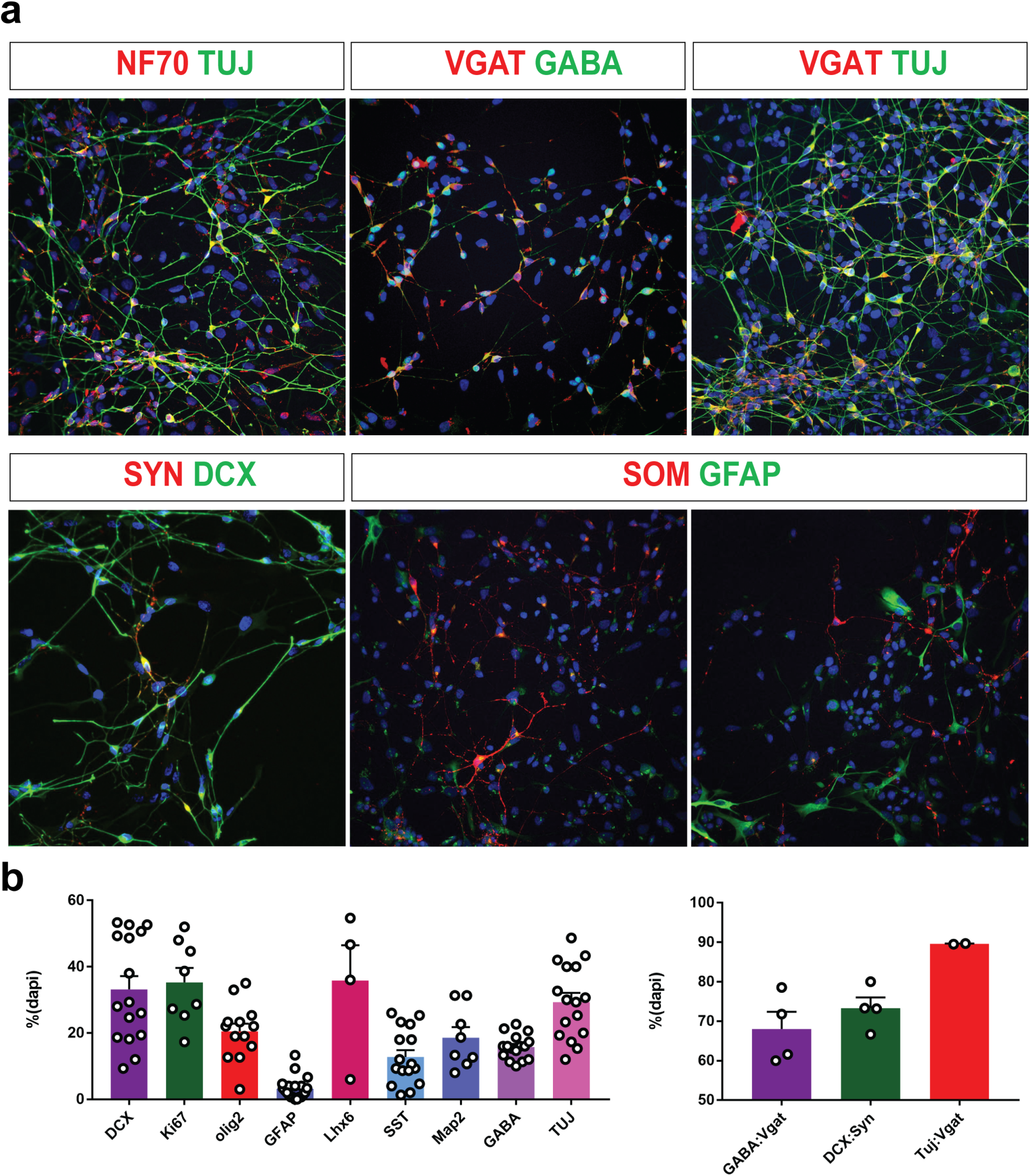
Migration of porcine embryonic MGE progenitors *in vitro*. (**a**) Matrigel explant assay showing migratory MGE progenitor cells (DAPI) immunolabeled with antibodies recognizing DCX^+^ and Lhx6^+^ cells (*top left*, merged image; *middle left*, individual images for DAPI, DCX and Lhx6; *bottom left*, higher magnification images for individual cells labeled with DCX and Lhx6; merged image at *far right*). Differential interference contrast bright-field image of embryonic explants at 4x magnification (top right). A homogeneous wave of migratory cells emerging from the explant core in all directions is marked in yellow; individual cells migrating to the outer reaches of the plate are marked in magenta (n = 12). (**b**) Schematics of the dissection approach for explant assay. Horizontal view of the GE (*top left*). MGE sub-dissected into 4 sections (AD, anterior-dorsal; AV, anterior-ventral; PD, posterior-dorsal; PV, posterior-ventral; *top middle*). Schematic representing from (a) showing distance measures from the explant (yellow and magenta arrows)(*top right*). Bar plot of explant migration assays measured as “distance from explant” (arbitrary unit) for all 4 MGE subregions (n = 4). Error bars represent s.e.m.

### *In vitro* differentiation of MGE progenitor cells

Under appropriate induction protocols (Franchi et al., 2018), MGE progenitors differentiate into pallial GABAergic neurons *in vitro*. Therefore, porcine MGE cells were cultured *in vitro* to evaluate molecular maturation. Cells dissected from E35 porcine MGE were plated onto polyornithine laminin coated dishes and cultured as monolayers. Immunohistochemical analysis was performed on day 14 (Fig. 7a). Many cells with mature neuronal morphologies and extensive processes co-expressed neurofilament (NF70) and neuron-specific class III β-tubulin (e.g., antigen for the Tuj1 antibody), a marker of immature and mature neurons. MGE-derived cells with complex neuronal morphologies expressed markers of inhibitory interneurons, namely GABA and a vesicular GABA transporter (VGAT). Consistent with an MGE identity, cells also expressed OLIG2 (20.5%), LHX6 (35.8%) and SOM (12.8%) (Figs. 7a-b). Many cells expressed DCX, a marker for young migratory neurons (Figs. 7a-b). Cells with neuronal morphologies also expressed synaptophysin (SYN), a synaptic vesicle protein and MAP2, a dendritic marker. Only a very small number of MGE-derived cells expressed GFAP (3.3%), an astrocytic marker. Undifferentiated stem cells were still present at this point in culture as indicated by expression of Ki67 (35.3%), a proliferative cell marker. The majority of interneuron-like GABA^+^ cells co-express VGAT (GABA:VGAT = 68%) or neuron-specific class III β-tubulin (TUJ:VGAT = 89.6%); significant co-expression of DCX and SYN was also noted (DCX:SYN = 73.3%)(Fig. 7c).

**Figure 7.**
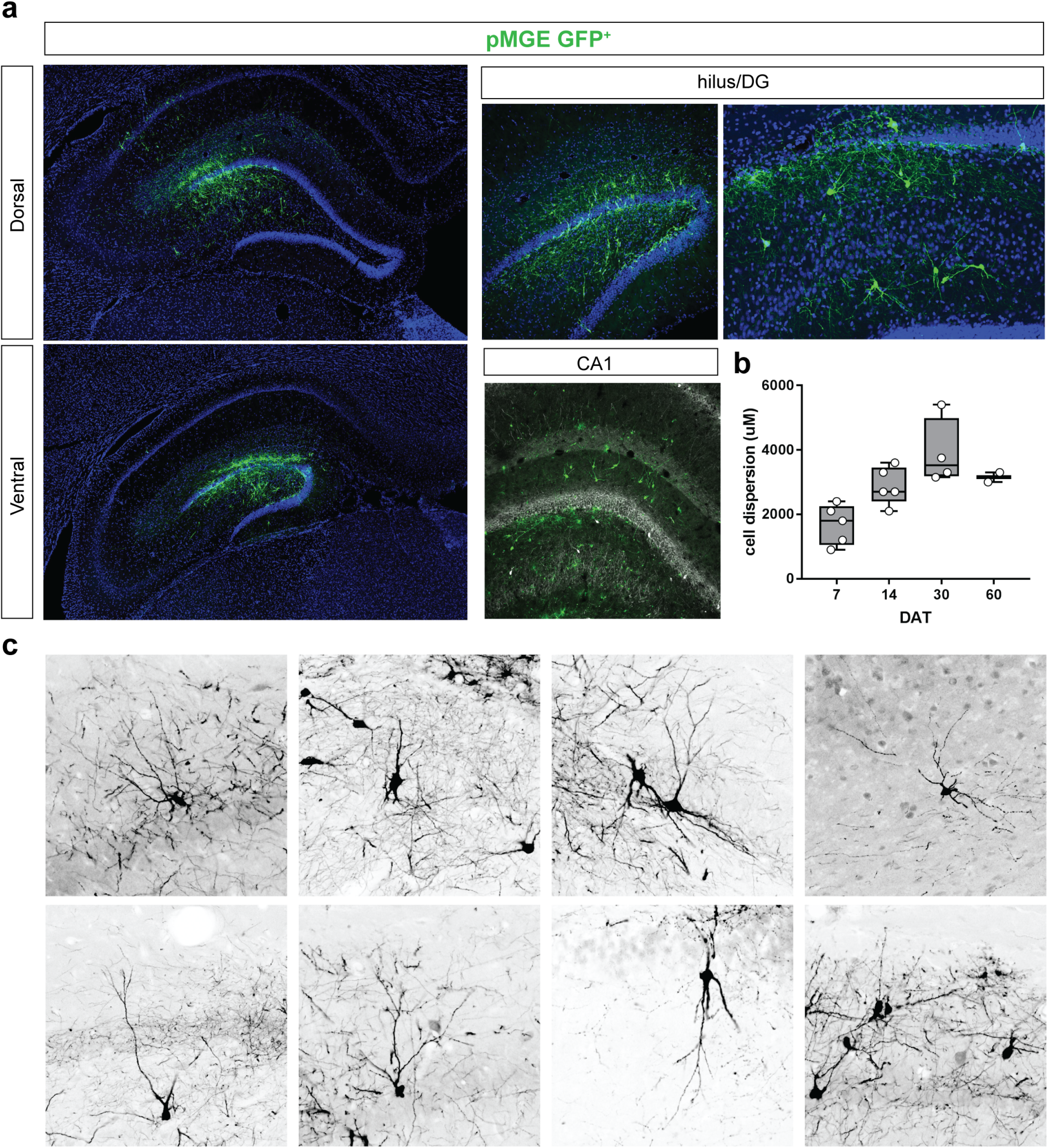
Characterization of porcine MGE progenitor cells *in vitro*. (**a**) Representative images of MGE-derived cells immunolabeled with markers of neuronal maturation (NF70, neurofilament; TUJ, neuron-specific class III beta-tubulin) and DCX, (doublecortin), interneurons (GABA, gamma-butyric acid; SOM, somatostatin or VGAT, vesicular glutamate transporter), astrocytes (GFAP, glial fibrillary acidic protein) or presynaptic terminals (SYN, synaptophysin). Images were collected at 14 days in culture. Quantification of expression plotted as a % over DAPI stain. Data are means ± s.e.m. (n = 3 or more independent experiments).

### Xenotransplantation of pig MGE in adult rats

Using embryonic brain slices (Anderson et al., 1997; Marin et al., 2000), *in utero* fate mapping (Wichterle et al., 2001), Matrigel explant assays (Wichterle et al., 1997, 2003), and *in vivo* transplantation (Wichterle et al., 1999; Alvarez-Dolado et al., 2006; Hunt et al., 2013), it is now well established that postnatal brain is permissive for migration and integration of transplanted embryonic murine MGE progenitors. To establish an efficient method for xenotransplantation and assessment of porcine MGE progenitors in a host brain, MGE was dissected from transgenic E35 pig embryos expressing GFP^52^. GFP expression was used to track migration and differentiation following transplantation into hippocampus of recipient adult wild-type Sprague-Dawley rats (n = 75; Table 1). At 30 DAT, GFP^+^ cells dispersed into most hippocampal sub-regions in dorsal and ventral directions with widespread distribution in dentate gyrus and CA1 (Fig. 8a-b). Distribution of porcine MGE-derived cells in host hippocampus increased between 7 and 14 DAT, reaching a relative plateau up to 60 DAT (Fig. 8c). Porcine MGE-derived GFP^+^ cells in hippocampus differentiated into cells resembling mature interneuron morphologies (e.g., bipolar cells, basket cells, and multi-polar cells) with elaborate dendritic trees and axonal projections (Figs. 8b, 8d). None of the porcine MGE-derived neurons exhibited morphological features of cortical pyramidal neurons (e.g., triangular cell soma extending a thick spiny apical dendrite). No tumors were observed in any host animal.

**Figure 8.**
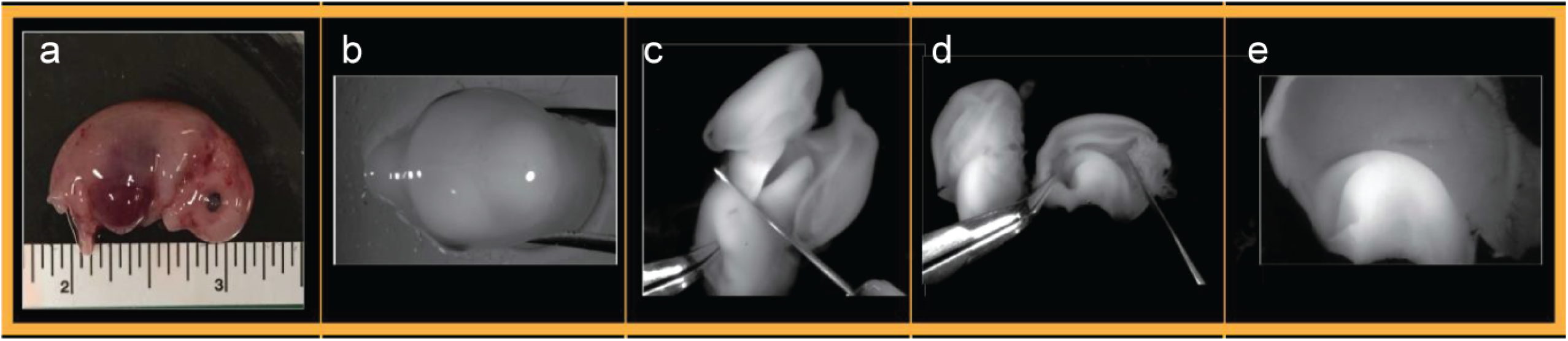
Distribution of porcine MGE-derived cells in the adult rat brain. (**a**) Hippocampus of adult Sprague-Dawley rat recipients 60 DAT labeled for DAPI (blue) and GFP^+^ transplanted porcine (p) MGE-derived neurons (green). MGE cells dispersed after xenotransplantation into recipient rats populating dorsal and ventral hippocampus locations. Higher magnification images show GFP^+^ cells in the rat hilus/dentate gyrus (DG) and CA1 subregions. (**b**) GFP^+^ cell dispersion along the anterior-to-posterior axis of the rat hippocampus at 7, 14, 30 and 60 DAT (n = 3 or more independent experiments). (d) Porcine MGE-derived cells in the rat hippocampus differentiated into neurons presenting typical morphology of interneuron subtypes e.g., bitufted, multipolar and basket.

**Figure 9.**
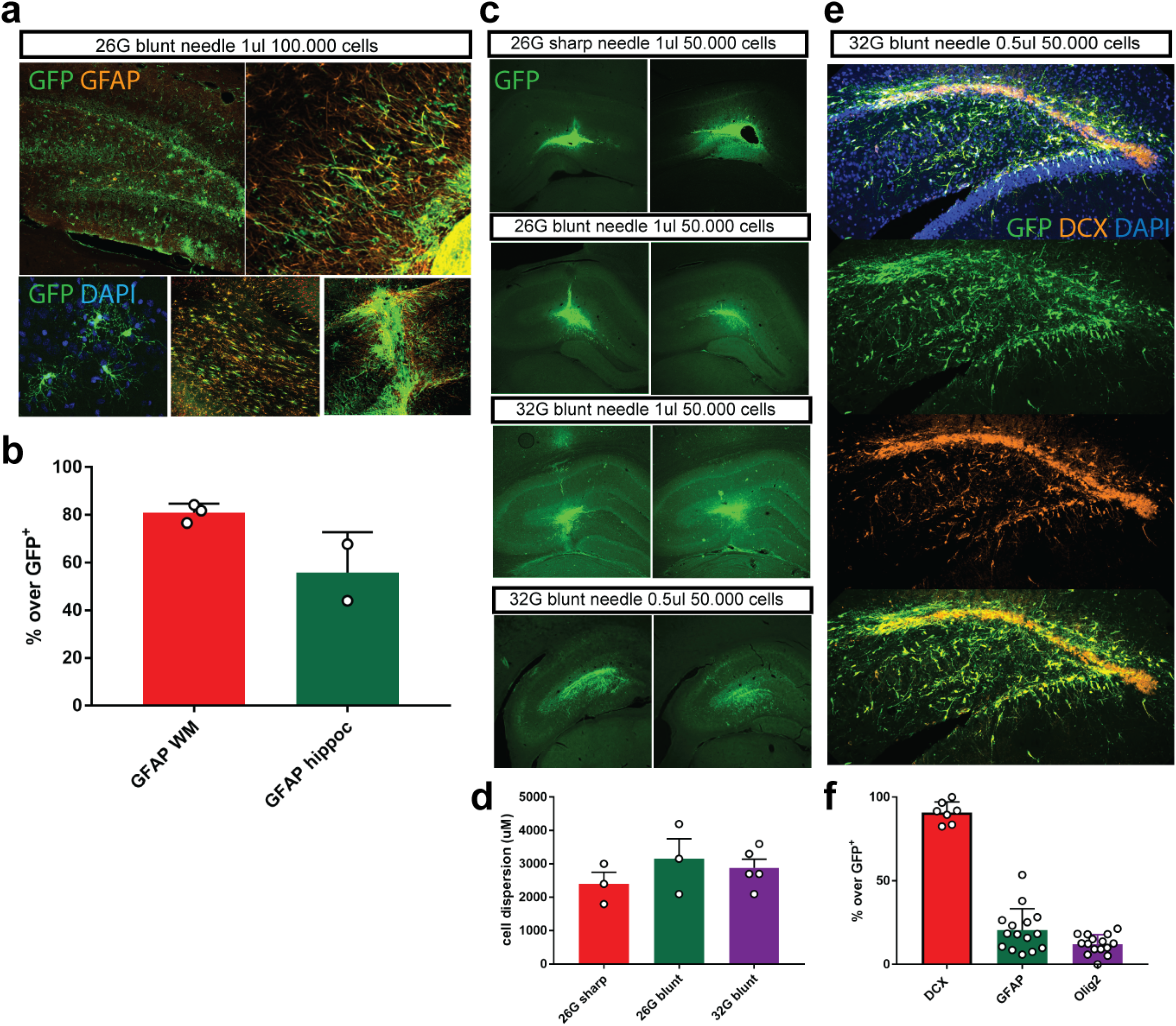
Dissection protocol for porcine MGE at E35. (**a**) Intact pig embryo. (**b**) Pig head viewed from the top. (**c**) Isolated pig brain with first posterior cut to isolate forebrain (**d**) Isolated pig brain micro-dissected into right and left hemispheres. Note: isolated right hemisphere has GE exposed. (**e**) Higher magnification image to view exposed GE region of right hemisphere.

**Figure 10.**
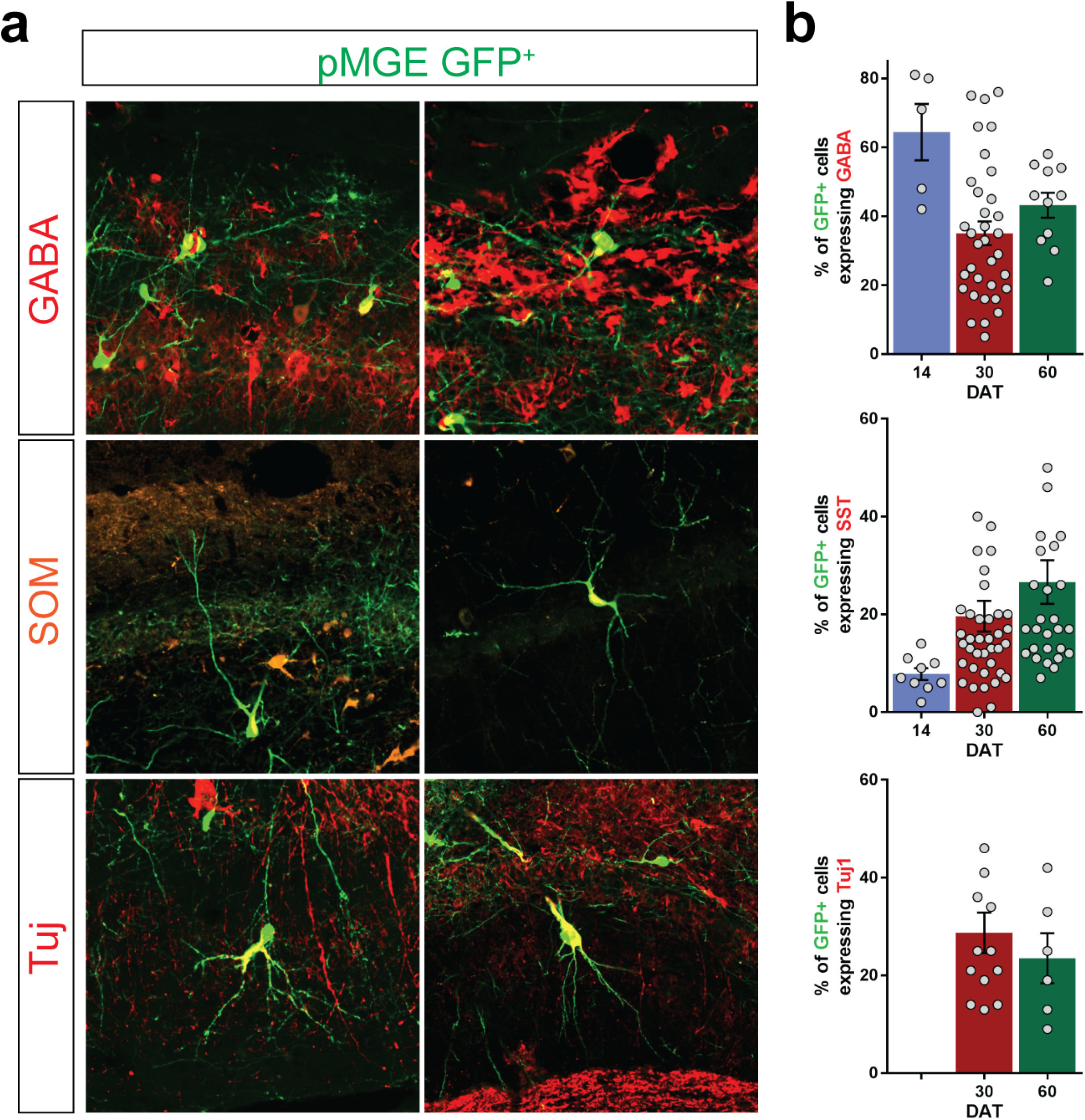
Optimization of porcine MGE xenotransplant protocol. A range of needle gauge sizes, injection volumes and cell densities were tested (n = 3 independent experiments per manipulation). (**a**) 100,000 porcine MGE GFP^+^ (green) cells in 1 μl transplanted into adult rat hippocampus using a 26G blunt Hamilton needle. Note the dense expression of GFAP-expressing MGE cells at 14 DAT (top panel). Higher magnification to show GFP^+^ cells with astrocytic morphology (lower left panel). Also note the high density of GFP:GFAP^+^ cells in fimbria and white matter tracts adjacent to hippocampus (lower middle panel). The injection site also shows a dense concentration of GFP^+^ cells unable to migrate out and astrocytes invading the needle tract (lower right panel). (**b**) Plot showing quantification of GFP:GFAP^+^ cells in white matter tract (80.9%) adjacent to hippocampus and inside hippocampus (55.9%). Mean ± s.e.m. (**c**) Representative images of the adult hippocampus (14 DAT) with GFP-labeled pMGE cells transplanted at varying injection needle gauges, injection volumes and cell densities. (**d**) Note the dense core of GFP^+^ cells near the injection site. Plot showing limited dispersion of GFP:GFAP^+^ cells in hippocampus. Mean ± s.e.m. (**e**) 50,000 porcine MGE GFP^+^ (green) cells in 0.5 μl transplanted into adult rat hippocampus using a 32G blunt Hamilton needle. Note the dispersion of GFP^+^ cells from the injection site and GFP:DCX^+^ neurons at 14 DAT (**f**) Plot showing % of GFP^+^ cells co-expressing DCX (90.9%), GFAP (20.5%), or Olig2 (12.1%).

MGE-derived interneurons can be identified by expression of the neurotransmitter GABA and neuropeptides such as SOM or PV (Fishell, 2008; Gelman et al., 2012; Laclef and Metin, 2018). Double-immunofluorescence revealed that most GFP^+^ porcine MGE-derived cells express GABA between 14 and 60 DAT. GFP^+^ cells exhibited increased subtype-specific SOM co-expression that also correlated with time post transplantation. By 60 DAT, the percentage of SOM:GFP^+^ cells significantly increased to 26.6% (Fig. 11). PV^+^ pMGE-derived neurons in the recipient rat brain could not be detected with available antibodies (see Methods). Tuj1 was present in GFP^+^ cells at 30 and 60 DAT. GFP^+^ cells expressing Olig2 (20.5%) were primarily localized to injection sites and only a small sub-population express GFAP (4.7%) (Fig. 12).

**Figure 11.**
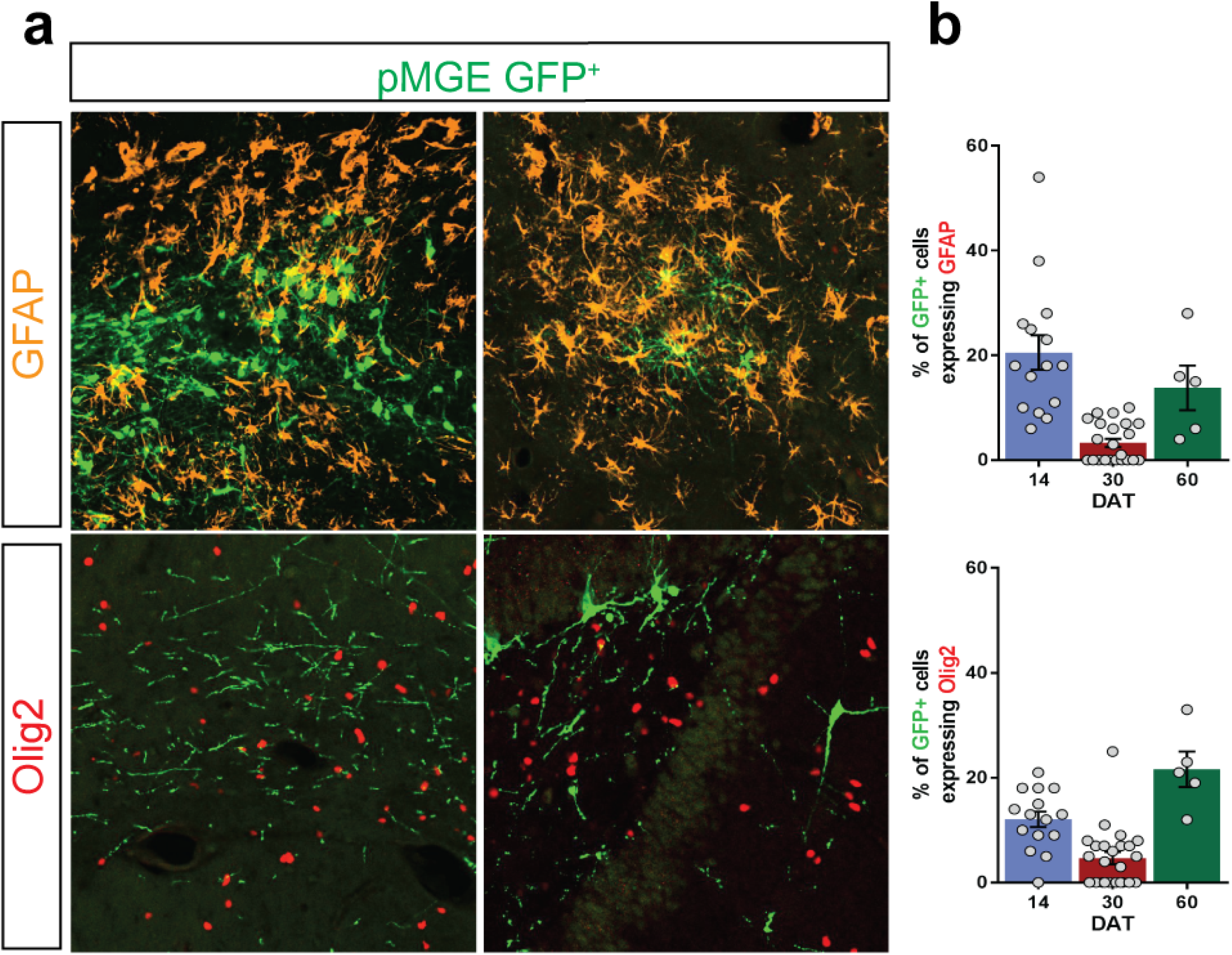
Immunohistochemical characterization of pMGE-derived cells in the adult rat brain. (**a**) Co-expression of GFP-labeled MGE-derived cells (green) with markers for inhibitory interneurons (GABA and SOM) or a neuronal marker (Tuj). (**b**) Quantification of pMGE-derived GFP^+^ cells co-expressing each marker at 14, 30 and 60 DAT. SOM 7 vs. 60 DAT: F(_2,32_) = 3.902; *P* = 0.0345, one-way ANOVA. Data are means ± s.e.m. (n = 3 or more independent experiments).

**Figure 12.** Immunohistochemical characterization of porcine MGE-derived cells in the adult rat brain. (**a**) Co-expression of GFP-labeled MGE-derived cells (green) with markers for astrocytes (GFAP) or oligodendrocytes (Olig2). (**b**) Quantification of MGE-derived GFP^+^ cells co-expressing each marker at 14, 30 and 60 DAT. Data are means ± s.e.m. (n = 3 or more independent experiments).

## Discussion

Here, we present strong evidence that porcine progenitors in embryonic medial ganglionic eminences express Nkx2.1, Lhx6, and Dlx2. *In vitro*, porcine MGE progenitor cells exhibit robust migratory properties and differentiate to neurons expressing GABA, vGAT and somatostatin. Following xenotransplantation into adult rat hippocampus, porcine MGE progenitors migrate extensively and differentiate to neurons with mature morphologies co-expressing GABA and somatostatin. Thus, porcine MGE progenitors may offer a valuable source of new inhibitory neurons for the types of transplantation-based therapeutic applications already suggested for mouse MGE progenitors e.g., epilepsy, Alzheimer’s disease and traumatic brain injury (Baraban et al., 2009; Anderson and Baraban, 2012; Hunt et al., 2013; Tong et al., 2014; Zhu et al., 2019).

The transcription factor profiles, migratory behavior (*in vitro* and *in vivo*) and differentiation into inhibitory interneurons observed in our porcine studies is remarkably similar to that described for mouse, monkey or human MGE. Seminal studies described a subpallial origin of cortical and hippocampal interneurons in rodents (Anderson et al., 1997; Marin et al., 2000; Pleasure et al., 2012), and a primary role of MGE in producing a subpopulation of interneurons has been extensively reviewed (Wonders and Anderson, 2006; Fishell, 2007; Gelman et al., 2012). From this work several families of transcription factors controlling subpallial embryonic development and fate of pallial interneurons originating in MGE are well established. These include Lhx6, Olig2, Dlx1/2 and Nkx2.1 in embryonic subpallial MGE (Du et al., 2008; Laclef and Metin, 2018) and Lhx6 in migrating MGE-derived cortical interneurons (Alifragis et al., 2004; Liodis et al., 2007; Vogt et al., 2014). It has also been demonstrated, through comparative analyses of gene expression patterns in chick, frog, zebrafish, lamprey, monkey and human, that basic developmental organization of the embryonic subpallium and pallial interneuron MGE origin is conserved across species (Hauptmann and Gerster, 2000; Puelles et al., 2000; Brox et al., 2004; Molnár et al., 2006; Cheung et al., 2007; Hansen et al., 2013; Pombal et al., 2011; Ma et al., 2013; Pauly et al., 2014). Our observations are entirely consistent with these descriptions.

In contrast to an earlier “failed” attempt to expand MGE-derived neurons *in vitro* from primary cells (Chen et al., 2013), our studies more closely replicate those of Franchi et al. (2018) showing that *in vitro* differentiation of rodent MGE progenitors into pallial interneurons involves outgrowth and elaboration of dendritic and axonal arbors (see Fig. 7). Similarly, most cells become GABAergic as indicated by co-expression of the inhibitory neurotransmitter GABA and presynaptic marker vGAT. At 14 DIV, porcine MGE cells also mimic the dense network of MAP2-positive dendrites and expression of a postsynaptic marker (synaptophysin) seen with rodent cultures. Unlike murine MGE, porcine MGE exhibited unique organization of DCX^+^ cells. This feature of porcine embryonic subpallium is reminiscent of human fetal brain sections (Hansen et al., 2013) and suggests a conservation of neurodevelopmental programs between pig and human not seen in rodents. It has been suggested that segregation into distinct cell clusters facilitates tangential migration of interneuron progenitors in developing brain. A process which, at the molecular level, may be regulated by expression of cell adhesion molecules such as fibronectin leucine rich-repeat transmembrane protein (Seiradake et al., 2014; del Toro et al., 2017) or transmembrane ephrin-B proteins (Zimmer et al., 2011; Dimidschstein et al., 2013). As such, further experimental analysis of porcine embryonic development could provide more direct insights into mechanisms underlying interneuron migration in higher species.

Another key attribute widely attributed to MGE progenitors is a capacity for migration (Wichterle et al., 1999). While most neuronal progenitors migrate through anatomically discontinuous cell-poor and cell-rich environments using radial glial scaffolds, chain migration allows migration of young GABAergic neurons through cell-dense regions independent of radial glial. Based on this unique migration capacity, MGE progenitors transplanted into cortex, hippocampus, striatum or even spinal cord, early in postnatal development or adult brain, retain a migratory capacity (Wichterle et al., 1999; Alvarez-Dolado et al., 2006; Baraban et al., 2009; Martinez-Cedeno et al., 2010; Braz et al., 2012; Hunt et al., 2013; Casalia et al., 2017). Taking advantage of these properties, MGE progenitor cell transplantation has shown remarkable therapeutic capacity in preclinical animal models of epilepsy (Baraban et al., 2009; Hunt et al., 2013; Casalia et al., 2017), traumatic brain injury (Zhu et al., 2014), Alzheimer’s disease (Tong et al., 2014), Parkinson’s disease (Martinez-Cedeno et al., 2010), and neuropathic pain (Braz et al., 2012; Juarez-Salinas et al., 2019). Here we used a xenotransplantation protocol to demonstrate that porcine MGE progenitors maintain this migratory capacity following transplantation into adult rat hippocampus. Robust migration was seen up to 60 DAT with cells differentiating to mature interneurons with complex dendritic arborizations and expression of antibodies marking MGE-specific inhibitory cell populations e.g., GABA and somatostatin. Only a relatively small number of cells expressed markers for astrocytes (GFAP) and unlike human ‘MGE-like’ stem cell derived interneurons (Carpentino et al., 2008) we never observed tumors. Previous work transplanting mouse MGE progenitors also consistently shows neurons expressing GABA and somatostatin (Alvarez-Dolado et al., 2006; Baraban et al., 2009; Hunt et al., 2013; Tong et al., 2014; Casalia et al., 2017; Zhu et al., 2019). While early postnatal MGE mouse transplant studies (and these vary by recipient brain region) report parvalbumin-positive neurons also consistent with an MGE lineage, PV^+^ cells take longer to develop and make up a smaller population of MGE-derived neurons in adult intra-hippocampal transplant studies (Hunt et al., 2013; Casalia et al., 2017). We also noted that unlike same species allograft murine-to-murine MGE transplant studies where immunosuppression protocols were unnecessary, even with a cocktail of immunosuppressants a pig-to-rat xenotransplantation protocol was only successful in about half of the animals. Not unexpected, as similar xenotransplant studies evaluating the fate of MGE-like human cells only utilized SCID immunodeficient mice as host animals (Nicholas et al., 2013; Maroof et al., 2014). The myriad immunoreactive processes that could take place, even in a relatively protected environment such as the central nervous system, remain to be optimized for these types of xenotransplantation studies. We anticipate that future studies will address these factors and perhaps utilize CRISPR-based gene editing technologies (Cowan et al., 2019; Hryhorowicz et al., 2020) to generate donor pig embryos ideally suited to this purpose.

## Acknowledgements

We thank Rosalia Paterno, Colleen Carpenter and Joseane Righes Marafiga for technical support and advice. We also acknowledge the National Swine Resource Center support in supplying pig embryos for our research under grant #U42-OD011140 (to NSRRC). This work was supported by an NIH R37 award (NS071785) from the NINDS and a UCSF Program for Breakthrough Biomedical Research award (to S.C.B.)

## Author contributions

M.F.C., M.L.P. and S.C.B. designed experiments. M.F.C., H.R., M.L.P. and T.L. carried out experiments and analyzed data. P.J.R. provided pig embryos and expertise. M.F.C. and S.C.B wrote the paper.

## Data availability

The data that support the findings of this study are available from the corresponding author upon reasonable request.

## Extended Data

**Time-lapse movie of lentivirus-labeled pig MGE.** Time-lapse movie showing migrating (green) MGE neurons in a cultured embryonic pMGE tissue slice at E35 (n = 3 experiments).

